# Improved humoral immunity and protection against influenza virus infection with a 3D porous biomaterial vaccine

**DOI:** 10.1101/2022.12.16.520784

**Authors:** Hiromi Miwa, Olivia Q Antao, Kindra M. Kelly-Scumpia, Sevana Baghdasarian, Daniel P. Mayer, Lily Shang, Gina M. Sanchez, Maani M Archang, Philip O. Scumpia, Jason S Weinstein, Dino Di Carlo

## Abstract

New vaccine platforms that properly activate humoral immunity and generate neutralizing antibodies are required to combat emerging and re-emerging pathogens, including influenza virus. Biomaterial scaffolds with macroscale porosity have demonstrated tremendous promise in regenerative medicine where they have been shown to allow immune cell infiltration and subsequent activation, but whether these types of materials can serve as an immunization platform is unknown. We developed an injectable immunization platform that uses a slurry of antigen-loaded hydrogel microparticles that anneal to form a porous scaffold with high surface area for antigen uptake by infiltrating immune cells as the biomaterial degrades to maximize humoral immunity. Antigen-loaded-microgels elicited a robust cellular humoral immune response, with increased CD4^+^ T follicular helper (Tfh) cells and prolonged germinal center (GC) B cells comparable to the commonly used adjuvant, aluminum hydroxide (Alum). By simply increasing the weight fraction of polymer material, we enhanced material stiffness and further increased antigen-specific antibody titers superior to Alum. Vaccinating mice with inactivated influenza virus loaded into this more highly crosslinked formulation elicited a strong antibody response and provided better protection against a high dose viral challenge than Alum. Thus, we demonstrate that by tuning physical and chemical properties alone, we can enhance adjuvanticity and promote humoral immunity and protection against a pathogen, leveraging two different types of antigenic material: individual protein antigen and inactivated virus. The flexibility of the platform may enable design of new vaccines to enhance innate and adaptive immune cell programming to generate and tune high affinity antibodies, a promising approach to generate long-lasting immunity against specific pathogens.

## Introduction

Despite tremendous advances in our understanding of immunology over the past decades, this knowledge has not translated into comparable advances in new vaccine material platforms for delivery of traditional protein/subunit or inactivated vaccines to extend and enhance the versatility of immunity generated. Successful vaccines have been developed against non-mutating pathogens like measles, poliovirus, and smallpox. However, the rapid evolution of mutating viruses including influenza virus and SARS-CoV-2 still results in huge socio-economic and public health challenges^[1,2]^. Like many RNA viruses, influenza has a much higher mutation rate than DNA viruses, making the development of a universal vaccine challenging^[3]^. Many vaccine strategies against influenza A virus have been implemented, including inactivated, live-attenuated and recombinant, to provide protection primarily by eliciting neutralizing antibodies^[4]^, however, these types of vaccines often fail to elicit adequate protection against an emergent strain with new mutations^[5]^. Contributing to the lack of long-term protection, antibody titers diminish after their peak at seroconversion, a few weeks after vaccination^[6–8]^.

To sustain antibody-mediated protection against a viral infection, a vaccine needs to elicit an antibody response against the correct epitope of the viral attachment protein that binds to host cells. To accomplish this, antigen-primed dendritic cells (DCs) from the immunization site migrate into secondary lymphoid organs where they prime CD4^+^ T follicular helper (Tfh) cells. These Tfh cells then instruct and select the best activated, antigen-specific B cells to undergo affinity maturation within germinal centers (GC)^[9–11]^. The formation of these GCs is critical for the evolution of antibodies to properly target neutralizing epitopes of key viral antigens and depends upon sustained antigen delivery ^[12–14]^. In addition to sustained antigen delivery, vaccine adjuvants often provide danger or other signals to prime immune responses to co-delivered antigens to recruit specific immune cell subpopulations and sustain the productive development of specific humoral or cellular immunity. There are currently only seven approved traditional vaccine adjuvant materials (including the newly approved Matrix M), many of which have limited control over the immune cell types that are activated, modest effects on activating these cells, and therefore do not promote a superior vaccine response for GC development, and these often poorly sustain antigen release ^[15,16]^. Ultimately, current adjuvant materials result in limited quantity, affinity, and breadth of GC-derived antibodies they can elicit, which negatively impacts the ability to neutralize current target antigens and future antigen variants following mutation. A new generation of highly tunable, immune-cell selective vaccine materials may help address these challenges.

Recent studies have reported that the physical properties of the vaccine delivery material are also critical to controlling the activity of vaccines^[17–20]^. When optimized, the antigen delivery material can prolong the time for antigen uptake, improve the bioaccumulation in lymphoid organs, effectively target specific immune cells based on the type of danger signal elicited, and overall elicit optimal immune responses. Polymeric microspheres and aluminum hydroxide, for example, have been widely investigated as vaccine carriers to provide sustained release^[21–23]^. This continuous release of antigen over time allows them to enhance the duration of the interaction between the antigen and the immune system^[24]^ and enhance the germinal center response and antibody levels^[14]^. Importantly, material stiffness cues in the microenvironment may directly polarize antigen-presenting cells (APCs) such as DCs involved in humoral immunity^[25–27]^. Stiffer substrates alter macrophage proinflammatory responses and polarize them towards an inflammatory phenotype^[28–30]^. Also, DCs grown in vitro on substrates with physiological stiffness were found to have reduced proliferation, activation, and cytokine production compared with cells grown on stiffer substrates^[31]^. Still vaccine materials have not taken full advantage of material physical properties, and are usually formulated with cytokines, growth factors, and pathogen associated molecular patterns to specifically recruit, target, and activate immune cells to elicit optimal immune responses^[32–34]^. The development of an antigen-releasing biomaterial that, through its physical properties, can sustain antigen delivery and provide proper immune activation signals for Tfh cells and GC B cell signals may overcome some of the supply chain, stability, and quality control challenges with these current approaches.

We have previously reported Microporous Annealed Particle (MAP) gels – an injectable biomaterial platform in which hydrogel microparticles are linked in situ to form a porous scaffold^[28,35–37]^. This hydrogel provides porosity at the scale of cells (tens of micrometers), preventing a foreign body response and fibrous encapsulation^[36,38,39]^. Recently, we found that treating skin wounds with a formulation of MAP where the chirality of amino acids within the crosslinking peptides was changed from L- to D-chirality (D-MAP) resulted in robust antibody responses to the antigenic D-amino acid-containing peptides and hair follicle regeneration in wounded skin^[28]^. Both recruitment of macrophages and hair follicle regeneration required an intact humoral immune response, suggesting that macroscale biomaterial scaffolds can directly influence humoral immunity and generate antibody responses to antigens contained within them^[40,41]^. Even in this antigenic formulation, the microporosity prevented fibrous encapsulation and foreign body responses, a desirable outcome for sustained delivery of an antigen^[42]^.

Given these features of the MAP biomaterial platform, we hypothesize that MAP can serve as a macroscale biomaterial-based vaccine platform for optimal antigen delivery that can be specifically tuned to activate the proper immune cells to achieve long-lasting and protective immunity against a viral pathogen without additional adjuvants. While 3D biomaterial scaffolds showed compatibility with mRNA vaccine platforms^[43,44]^ and as cancer vaccines^[45]^, the role of a 3D porosity for traditional (recombinant antigen/protein and inactivated virus/particle) vaccines has not been investigated in detail. Herein, we test whether we can create vaccine-MAP scaffolds (VaxMAP), in which antigens or inactivated viral particles are embedded in the microgels during fabrication. We found that the VaxMAP platform activated an adaptive immune response and generated protective antigen-specific antibodies against a model antigen and inactivated influenza particles. By simply increasing the weight percentage of PEG within the microgels, which resulted in a more heavily cross-linked, and therefore stiffer hydrogel, we induced greater Tfh cell expansion, prolonged GC B cell development, and greatly enhanced high affinity antibody responses. The inactivated influenza virus particles (iIVP)-MAP elicited antibody responses against hemagglutinin and protected against a lethal influenza virus infection. Thus, VaxMAP represents a tunable, injectable biomaterial platform to enhance humoral immunity for traditional vaccines.

## Results and Discussion

Microfluidic emulsion generation-based polymerization allows the manufacture of highly customizable microparticles **(Figure 1A and Figure S1, Supporting Information)**. VaxMAP is composed of matrix metalloproteinase (MMP) sensitive polyethylene glycol (PEG)-based microgel beads decorated with K and Q peptides that are substrates for transglutaminases and allow the linkage of the microgels to each other and surrounding tissue. Monodispersed VaxMAP microgels (77.1±4.6 μm diameter average) were produced using a microfluidic droplet generator with consistent swelling ratios from particles with a total diameter ranging from 50 to 110 μm (**Figure 2A**). The concentration of encapsulated antigen was controlled by changing the initial antigen concentration within pre-gel solutions **(Figure 2B)**. We also examined the influence of PEG concentration on the mechanical stiffness of MAP gels to investigate potential effects on immune modulation *in vivo*^[46,47]^. By increasing the PEG concentration from 5% to 7.5% the microgels had a higher crosslinking density and an increased hydrogel stiffness, resulting in a storage modulus that increased from ∼1000 Pa to ∼4000 Pa **(Figure 2C)**. A scaffold of packed microgel particles also resulted in cell-sized interconnected voids that led to higher convective flux of fluid (**Figure 2D, E**). Despite only limited permeation occurring in the nonporous bulk hydrogel, the interconnected porosity of annealed MAP hydrogels derived from both 5% and 7.5% PEG microgels resulted in a more than 100-fold enhancement in fluid conductivity **(Figure 2F and Figure S2, Supporting Information)**. In order to assess the generation of a humoral response and antibody affinity, w e incorporated a classical immunological antigen, 4-hydroxy-3-nitrophenylacetyl hapten conjugated to ovalbumin (NP-OVA) antigen (MW = 45kD) into the gel matrix at various concentrations during manufacture, without affecting the manufacturing process and found the NP-OVA would remain entrapped over at least a 3-day period (**Figure 2G**). We also found that upon exposure to degradative enzymes *in vitro*, the NP-OVA was slowly released as the microgel particles swelled (**Figure 2G**). In addition, *in vitro* culture of macrophages and dendritic cells with fluorescently-conjugated MAP gel resulted in fluorescently-labeled components accumulating within the cells after 1 day of culture and increasing over 7 days of culture with macrophages being able to take up more fluorescent label than DCs following breakdown of the material (**Figure 2H and Figure S3, Supporting Information**).

**Figure 1.**
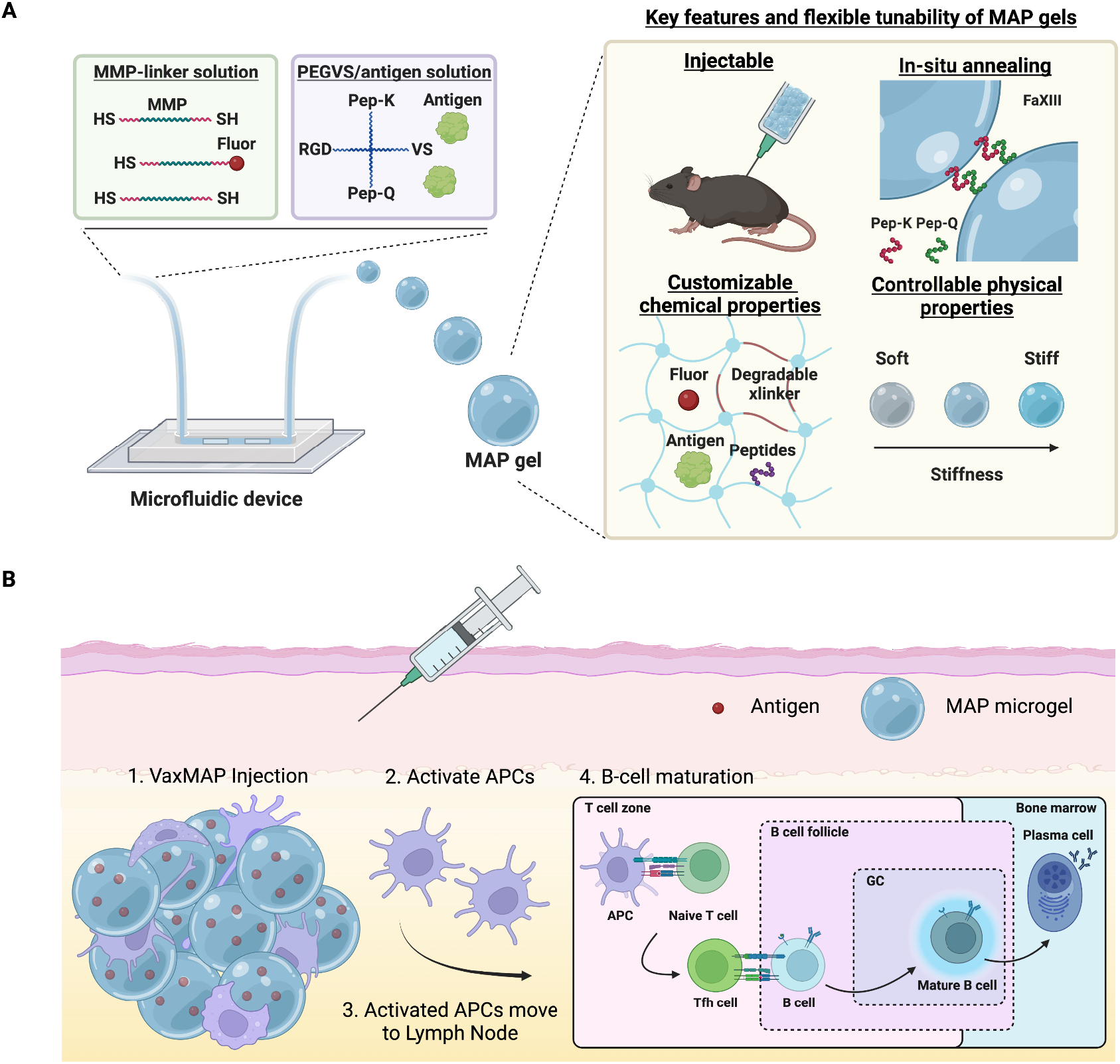
Scheme illustrating the microfluidic generation of VaxMAP scaffolds for vaccination. A) MAP microgels are manufactured using microfluidic droplet emulsion polymerization from pre-gel solutions containing 4-arm PEG-vinylsulfone (PEG-VS) via thiol-ene reactions to encapsulate antigen in the dense gel mesh. Matrix metalloproteinase (MMP) degradable peptide linkers enable gel degradation. RGD peptides are present on the microgels to facilitate cell infiltration into the void spaces between gels, maximizing surface area for cell-biomaterial contact. The MAP microgel slurry is easily injectable and K and Q peptides enable linkage of microgels to tissue and each other by activated Factor XIII once injected. Microgel stiffness is controlled independently of cell-scale porosity. B) Pores between spherical gels allow immediate cellular infiltration. Gels are degraded and antigen is uptaken by antigen presenting cells (APCs), which then direct an adaptive immune response. Tfh cell development is initiated in the T cell zone where naïve T cell differentiation is instructed by antigen-primed dendritic cells. Newly formed Tfh cells and antigen-activated B cells meet at the border of the T cell zone and B cell follicle and following productive interactions, enter the follicle to form a germinal center (GC). Within the GCs, B cells undergo selection and differentiate into plasma cells.

**Figure 2.**
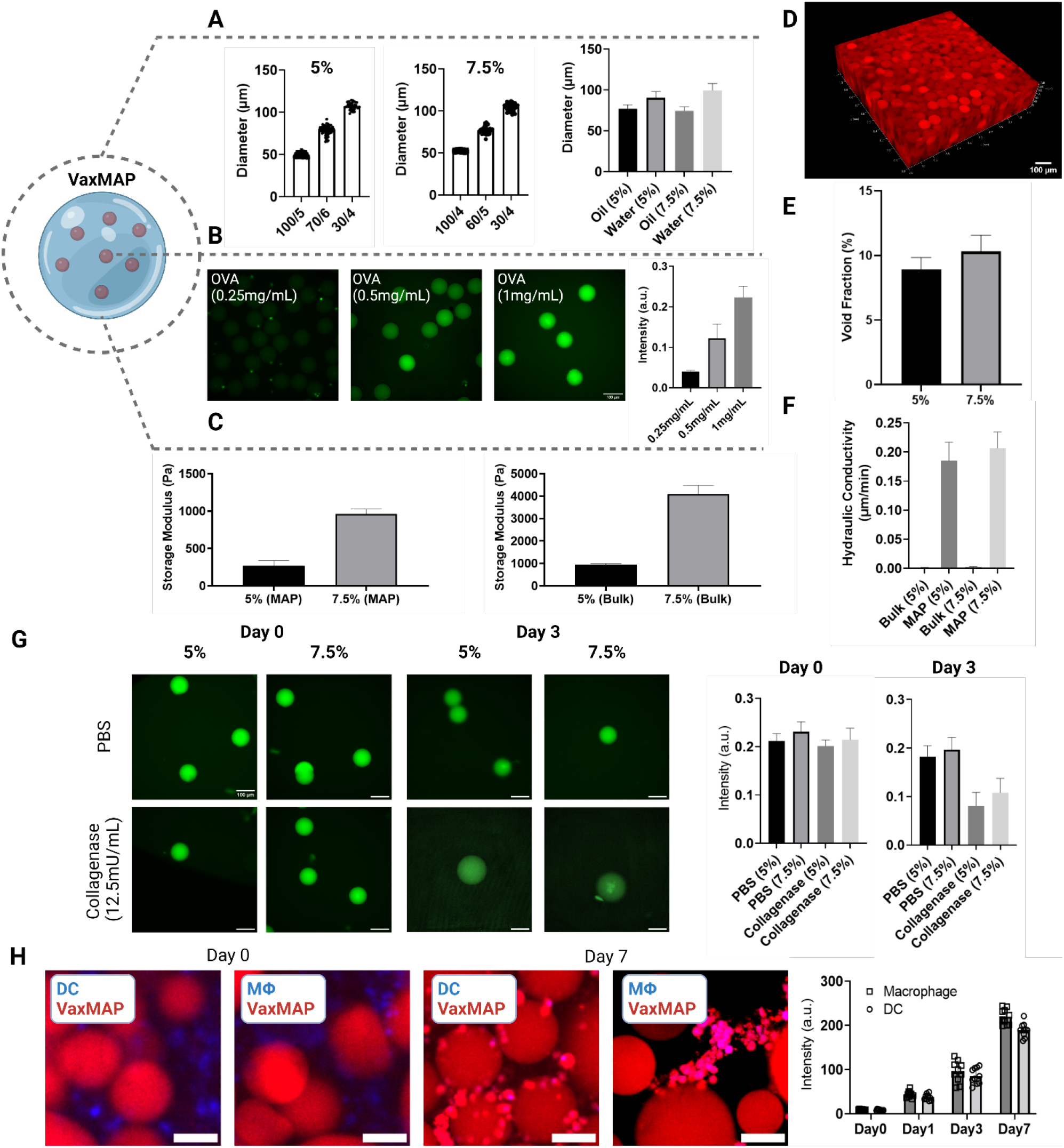
Characterization of VaxMAP building blocks and annealed scaffolds. A) Generation of VaxMAP building blocks with highly defined sizes by altering the aqueous flow rate. Particle size distributions are shown for different ratios of oil to pre-polymer for 5% and 7.5% PEG microgels. VaxMAP building blocks were made with an aqueous flow rate of 6 or 5 μL min^−1^ for 5% and 7.5% PEG gels respectively, and swollen in buffer after aqueous extraction from the oil phase. B) Representative images of VaxMAP gels loaded with increasing amounts of fluorescently-labeled OVA. With increasing OVA concentration, the fluorescence signal correspondingly increased. Scale bar: 100 μm C) Stiffness of VaxMAP hydrogels formed in bulk and annealed MAP gel form. Polymer concentration contributes to an increment in stiffness. D) Fluorescence confocal image of a scaffold assembled from monodisperse fluorescently-labeled MAP building blocks. Scale bar: 100 μm. E) Void fraction of annealed MAP scaffolds. F) Hydraulic conductivity of PBS through a nonporous bulk scaffold and annealed MAP scaffold at atmospheric pressure. G) Fluorescent OVA antigen is imaged in microgels over 3 days. Average microgel intensity remains stable in PBS but decreases with collagenase treatment as microgels also swell in size. The scale bar is 100 μm. H) Mouse bone marrow-derived primary macrophages and dendritic cells co-cultured with Alexa Fluor 555-conjugated MAP scaffolds. Over 7 days of culture cells uptake MAP components and cell-associated Alexa Fluor 555 signal increased. Blue: DAPI stain, Red: Alexa Fluor 555. The scale bar is 100 μm. Data are shown as mean ± SD.

We found that MAP gels injected and annealed *in vivo* in mice led to activation of CD4^+^ T cells and B cells resulting in the generation of antigen-specific antibodies, a hallmark of an effective vaccine response. We incorporated NP-OVA as described above **(Figure 2G)**, into MAP to characterize T and B cell responses, including high affinity antigen specific immune responses. C57Bl/6 mice were immunized subcutaneously (s.c.) with MAP microgels (5% PEG) loaded with NP-OVA (MAP-NP-OVA), or the standard aluminum hydroxide-based adjuvant, Alum loaded with NP-OVA (Alum-NP-OVA)^[48]^ and compared to unimmunized, PBS injected mice, where activated T and B cells in lymph nodes were assessed 12 days later. MAP-NP-OVA immunization induced similar percentages of activated CD4^+^ T cells (CD4^+^CD44^hi^) and B cells (B220^+^IgD^lo^) compared to Alum-NP-OVA **(Figure 3B and 3C)**. To assess if MAP immunization induces NP-specific antibody responses, we performed anti-NP ELISAs from sera collected at day 12 after immunization. Antigen-specific IgG1 antibodies increased following immunization in C57Bl/6 mice, antibodies in mice class-switched to IgG1 correlate with type 2 immune polarization (Th2-biased) consistent with the expected MAP and Alum priming^[28,49]^. MAP-NP-OVA immunized mice displayed modestly higher titers of total anti-NP IgG1 compared to Alum-NP-OVA whereas PBS controls did not display detectable anti-NP antibodies **(Figure 3D)**. Since the immune response following alum immunization typically resolves by 28 days^[50]^, we examined whether the anti-NP antibody responses remained elevated 28 days following immunization with MAP-NP-OVA, Alum-NP-OVA or PBS. Total anti-NP specific antibody titers remained modestly elevated in MAP-NP-OVA immunized mice compared to Alum-NP-OVA **(Figure 3E)**. Neither MAP-NP-OVA or Alum-NP-OVA induced significant anti-NP specific IgG2c responses, consistent with Th2 polarization of antibody responses (**Figure 3F and G**). In concert, these data demonstrate that vaccination with MAP-NP-OVA induces T and B cell activation and anti-NP antibody responses as well an Alum-NP-OVA immunization. Thus, immunization with MAP activates B and T cells to generate an antibody response towards delivered antigens.

**Figure 3.**
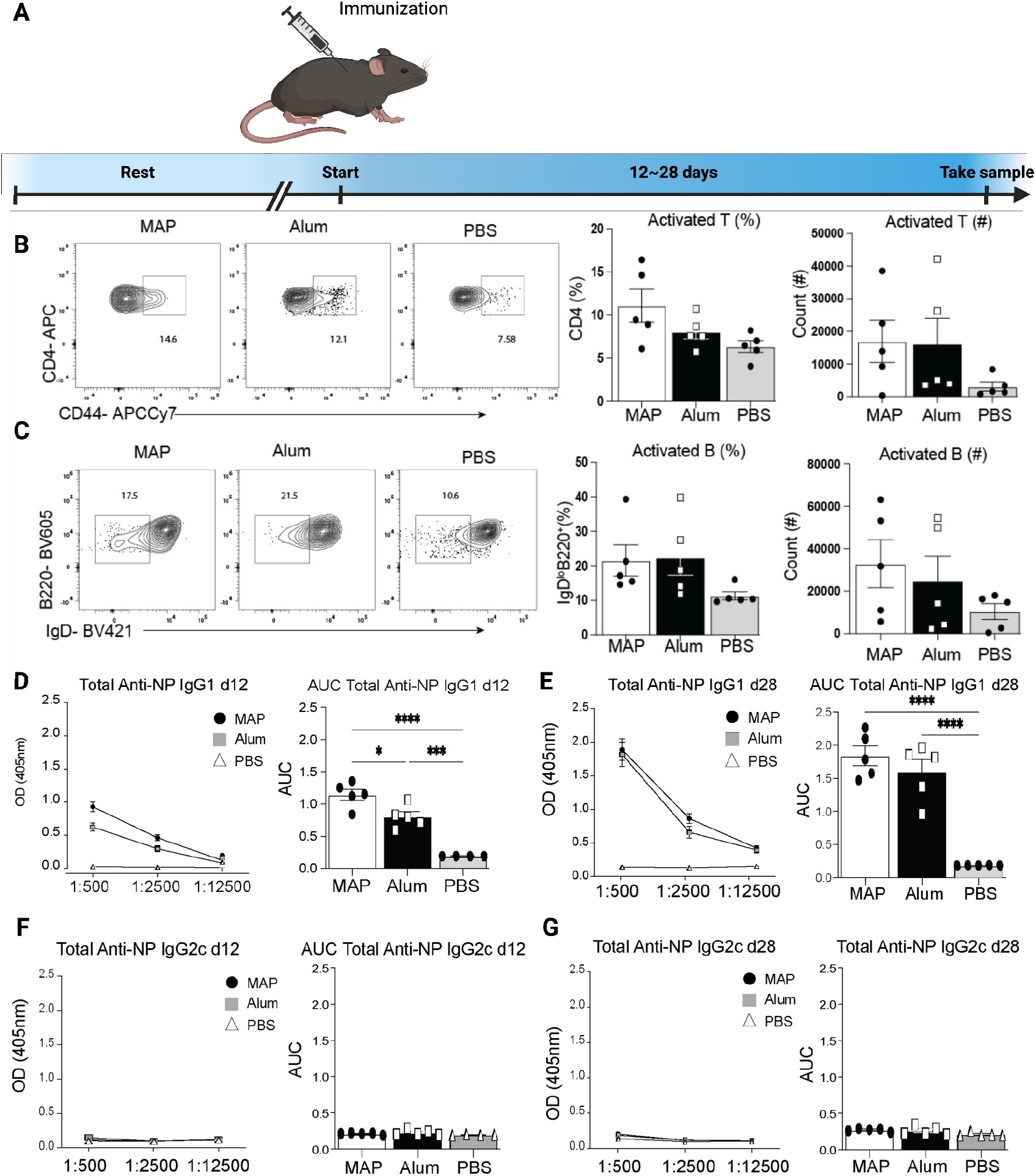
MAP vaccination induces robust T cell and B cell activation and anti-NP antibody production. A) Experimental timeline. B) Representative FACS plots and graphs of percentages and counts of activated CD4+ T cells. C) Representative FACS plots and graphs of percentages and counts of activated B cells. D) Optical Density and Area Under the Curve of anti-NP IgG1 antibodies at 12 days post immunization. E) Optical density and area under the curve of anti-NP IgG1 antibodies at day 28 post immunization. Optical density and area under the curve of anti-NP IgG2c antibodies at day 12 or 28 post immunization (F) and (G), respectively.

We next assessed whether the high affinity antigen-specific antibody responses driven by MAP-NP-OVA immunization were associated with increased numbers of activated Tfh and GC B cells. To examine if Tfh and GC B cells develop following MAP injection, C57Bl/6 mice were immunized s.c. with MAP-NP-OVA, Alum-NP-OVA, and PBS and assessed 12 days later. Indeed, the percentage and numbers of Tfh cells in draining lymph nodes (LN) trended higher in MAP-NP-OVA immunized mice compared to Alum and PBS injected mice **(Figure 4A)**. In accordance with elevated Tfh cells, the percentage of GC B cells were increased in MAP-OVA immunized mice compared to Alum and PBS injected mice **(Figure 4B)**. Furthermore, consistent with slower antigen release, Tfh and GC B cells remain elevated at 28 days post immunization with MAP-NP-OVA compared to Alum and PBS immunized mice (**Figure S4, Supporting Information**). To assess whether the increase in Tfh and GC B cells observed in MAP-NP-OVA injected mice resulted in elevated high-affinity antigen-specific antibodies, we measured the levels of high affinity anti-NP IgG1 antibodies from immunized mice at days 12 and 28. While high affinity antibodies were lower in MAP-NP-OVA immunized mice compared to Alum at day 12 **(Figure 4C)**, the former had elevated titers of high affinity anti-NP IgG1 antibodies by day 28 post immunization **(Figure 4D)**, consistent with a prolonged GC response. Similar to their type 2 immune responses and generation total antigen-specific antibodies either immunization induced significant anti-NP specific IgG2c responses, Thus, VaxMAP generated a prolonged immune response compared to a traditional vaccine platform.

**Figure 4.**
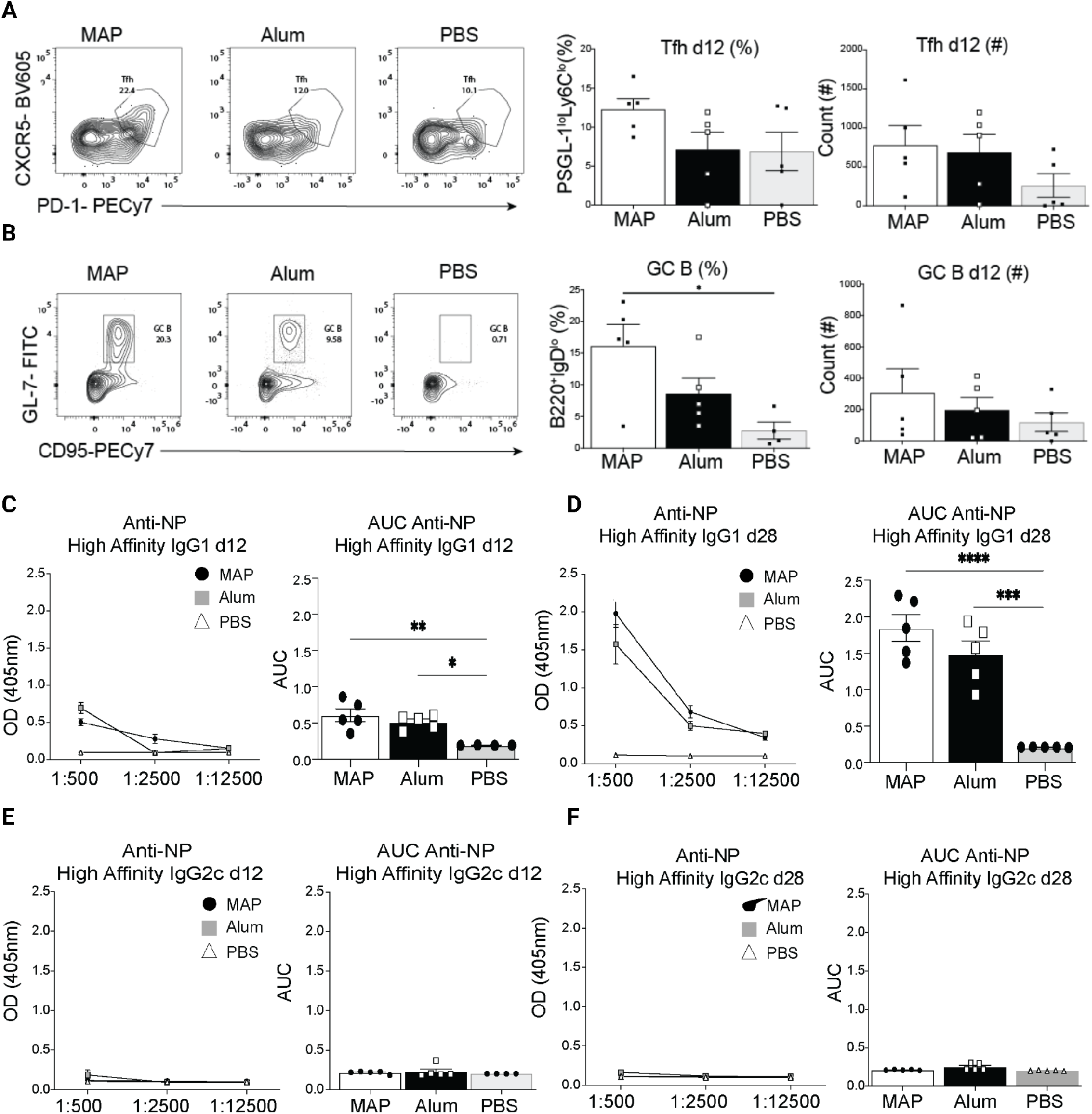
MAP vaccination induces Tfh and GC B cell development and production of high affinity antibodies. A) Representative FACS plots, percentages and counts of Tfh cells at day 12 post immunization. B) Representative FACS plots, percentages and counts of GC B cells at day 12 post immunization. C) Optical density and area under the curve for high affinity anti-NP IgG1 antibodies at day 12 post immunization. D) Optical density and area under the curve of high affinity IgG1 antibodies at day 28 post immunization. Optical density and area under the curve of high affinity anti-NP IgG2c antibodies at day 12 or 28 post immunization (E) and (F), respectively.

Since MAP fabrication is done under aseptic techniques but is difficult to keep completely sterile throughout the manufacturing process, we next confirmed that endotoxin contamination did not contribute to the immune response elicited by VaxMAP as TLR4-mutant C3H/HeJ mice had similar antigen-specific antibody response (Data not shown). We next examined whether MAP elicited innate immune cell activation through any traditional pathogen- or danger-associated molecular pattern (PAMP or DAMP, respectively) pathways. Since activation of all TLR and inflammasome-interleukin (IL) 1 receptor (IL-1R) pathways require signaling through the adaptors Myeloid differentiation primary response 88 (MyD88) and/or Toll-IL-1R domain (TIR)-containing adapter-inducing interferon-β (Trif), we immunized B6 and *Myd88*^*-/-*^*Trif*^*-/-*^ double knockout (DKO) mice with MAP-NP-OVA. We found that similar to TLR4 mutant HeJ mice, DKO mice displayed no reduction in antigen specific IgG1 antibodies at Day 14 and Day 28 **(Figure S5, Supporting Information)**. Overall, these data illustrate that MAP promotes humoral immune responses to antigens delivered within the microparticles that is not impacted by indirect endotoxin contamination or traditional PAMP/DAMP signals, but instead rely on direct interactions with the antigen present within the biomaterial to generate a type 2 antibody (IgG1) response to the antigen.

Since the antibody responses to immunization with antigens in MAP did not seem to rely on traditional PAMP/DAMP signaling pathways, we next wished to examine whether modifying the physicochemical properties of the particles could result in enhanced immune responses. Since biomaterial/matrix stiffness participates in immune cell activation and polarization through enhanced mechanotransduction in innate immune cells^[25,26,51]^, we next wished to determine whether enhancing the stiffness of the hydrogel further could further enhance the antibody response. Since increased cross-linking density can result in increased stiffness while simultaneously allowing for slower release of proteins from hydrogel^[52]^, we chose to modulate PEG crosslinking density to enhance adjuvant effects of MAP. We generated two formulations of MAP-NP-OVA, one formulation using 7.5% PEG and the same MMP-degradable cross-linking peptide, and another with 7.5% PEG cross-linked with a PEG-dithiol crosslinker instead. Both formulations resulted in increased stiffness when compared to the 5% MAP formulation, with the 7.5% MAP with dithiol crosslinker resulting in the highest stiffness **(Figure S7, Supporting Information)**. We found that both total and high affinity anti-NP specific antibody production were enhanced by both formulations of 7.5% MAP over 5% MAP, with a slight enhancement of dithiol crosslinked MAP over peptide crosslinked MAP **(Figure S7, Supporting Information)**. While the PEG crosslinked hydrogel led to slightly better antibody responses, three of the mice developed scratching behavior and ulcerations over the injection site, either due to the uncomfortable nature of the dithiol crosslinked stiffer hydrogel or possibly due to the more intense immune response in the area. Given their ability to induce a nearly maximal antigen-specific antibody production without causing any obvious undesired side effects, the 7.5% MMP crosslinked gels were chosen as the preferred formulation for future studies.

To examine how more highly crosslinked MAP formulations regulate the immune response, we first immunized C57Bl/6 mice with either 5% or 7.5% MAP-NP-OVA and assessed T and B cells and antibody responses 12 days post injection. The stiffer 7.5% MAP gel induced enhanced CD4^+^ T cell activation as well as Tfh cell development compared to the 5% MAP formulation **(Figures 5A and B)**. Despite the 7.5% MAP gel having reduced activated B cell percentages **(Figures 5C)**, GC B cells were increased compared to the 5% MAP **(Figures 5D)**. We next compared if the increased Tfh and GC B cells response in the 7.5% MAP-NP-OVA also resulted in improved antibody responses of mice immunized with NP-OVA in the traditional Alum adjuvant. Indeed, the 7.5% NP-OVA-MAP formulation induced elevated anti-NP IgG1 antibody responses compared to 5% MAP- or Alum-based immunizations **(Figures 5E)**, indicating that the 7.5% MAP formulation drove a more robust antibody response than Alum. Importantly, while the 7.5% MAP-NP-OVA formulation elicited more robust anti-NP IgG1 responses, it did not change the polarization of the immune response, as anti-NP IgG2c responses remained low (**Figure S7, Supporting Information)**. Thus, changing the crosslinking density of a biomaterial can result in enhanced immune responses without altering the polarization of the immune response and represents a novel way to enhance antigen-specific immunity.

**Figure 5.**
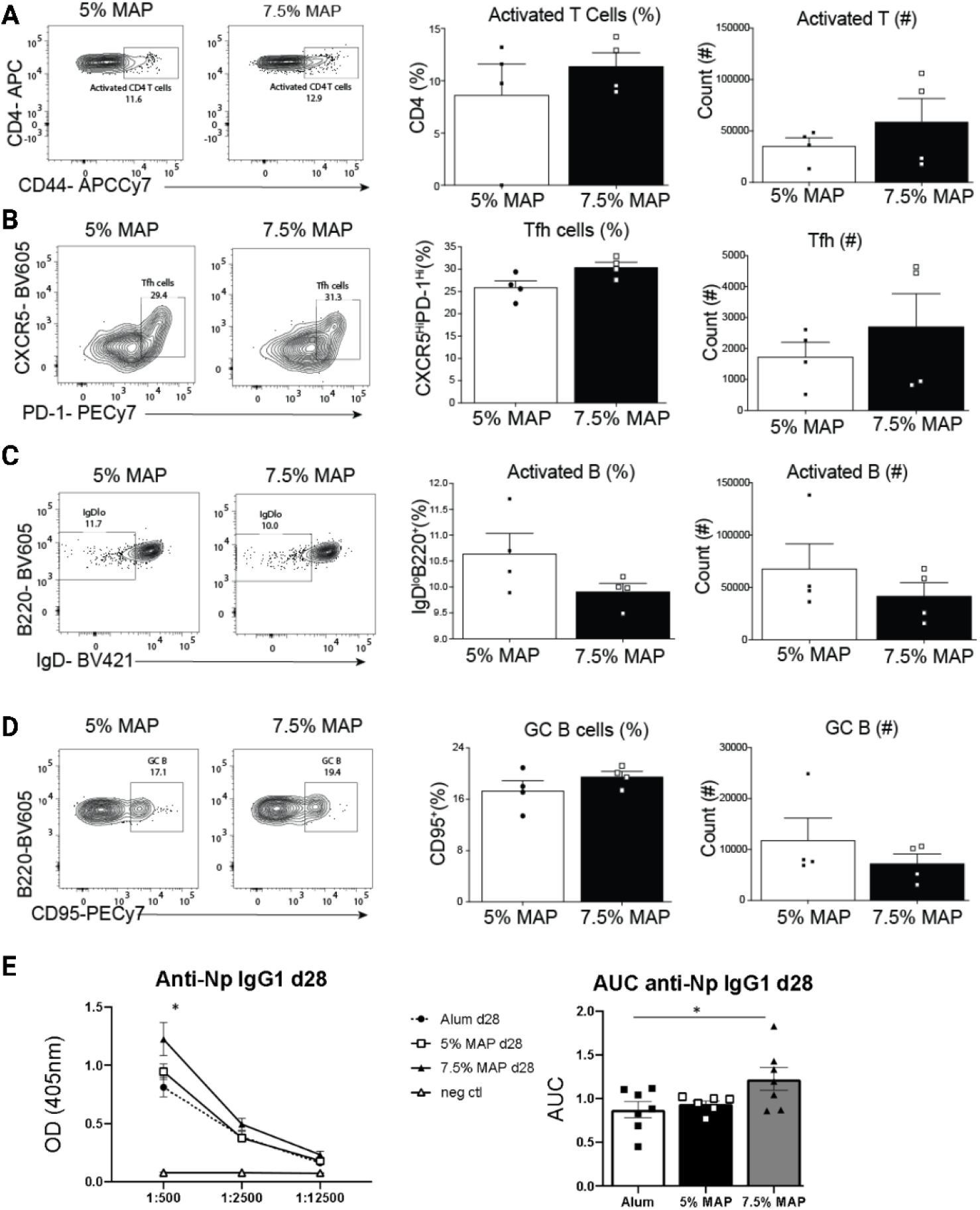
Effect of VaxMAP crosslinking density on the adaptive immune response 12 days after immunization. A) Representative FACS plots, percentages and counts of activated T cells between 5% and 7.5% MAP formulations. B) Representative FACS plots, percentages and counts of activated B cells between 5% and 7.5% MAP formulations. C) Representative FACS plots, percentages and counts of Tfh cells between 5% and 7.5% MAP formulations. D) Representative FACS plots, percentages and counts of GC B cells between 5% and 7.5% MAP formulations. E) Optical density and area under the curve of high affinity anti-NP IgG1 antibodies of mice immunized with 5%, 7.5% MAP, Alum and the unimmunized PBS control.

While NP-OVA is an advantageous system for dissecting immune responses, it is not naturally expressed by pathogens and does not reflect the ability to confer protection against infection. Using the 7.5% MAP formulation, which led to increased antibody titers, we next evaluated the ability to introduce whole inactivated virus from the mouse-adapted influenza strain (PR8) into MAP (MAP-flu), compared to the same particles delivered in Alum (Alum-flu) or PBS (PBS-flu). C57Bl/6 mice were immunized with MAP-flu, Alum-flu, or PBS-flu as a control and immune responses were examined 12 days following injection (Figure 6A). We found that MAP-flu drove the development of activated T cells greater than PBS controls **(Figure 6B)**.

**Figure 6.**
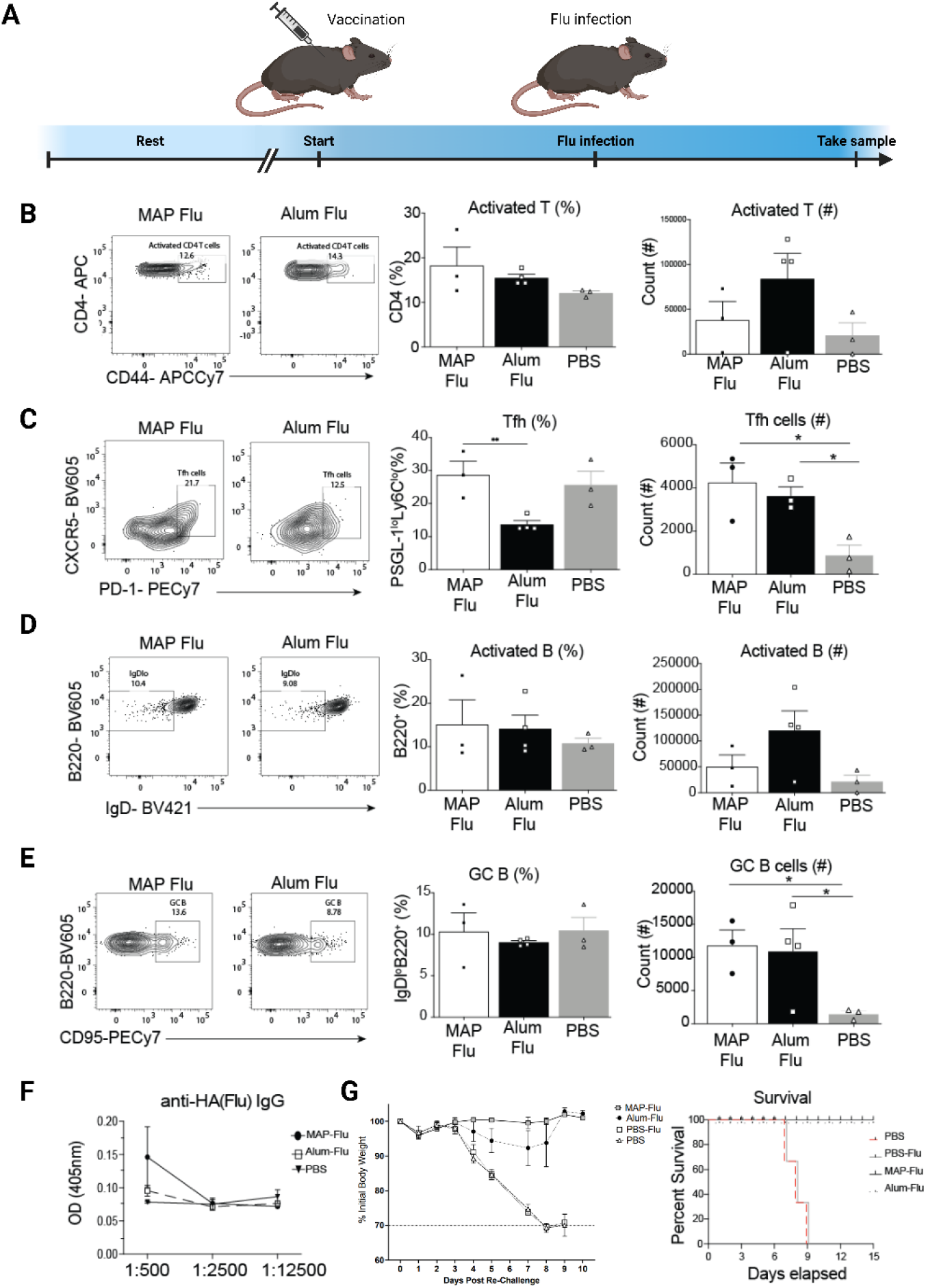
MAP is an effective vaccination platform against PR8 influenza infectionn. A) Experimental timeline. B) Representative FACS plots, percentages and counts of activated T cells of immunized mice. C) Representative FACS plots, percentages and counts of Tfh cells of immunized mice. D) Representative FACS plots, percentages and counts of activated B cells. E) Representative FACS plots, percentages and counts of GC B cells. F) Optical density and area under the curve of anti HA IgG antibodies. G) Daily percent of individual mouse weight loss from each group and survival curve following PR8 influenza rechallenge 60 days following immunization.

Additionally, MAP-flu induced a significantly higher percentage of Tfh cells than Alum-flu or PBS-flu immunized mice **(Figure 6C)**. In a similar manner, MAP-flu immunization induced modest activation of B cells and GC B cell development **(Figures 6D and E)**. Anti-hemagglutinin (HA) IgG antibody responses were increased in mice receiving MAP-flu compared to PBS immunized controls **(Figure 6F)**. To assess if MAP-flu immunization elicits protection against influenza infection, C57Bl/6 mice were immunized with either MAP-flu, Alum-flu, PBS-flu, or PBS alone. Sixty days later, these mice were challenged with a high dose of PR8 influenza. Mice immunized with MAP-flu demonstrated enhanced survival relative to PBS-flu and PBS immunized mice **(Figure 6G)**. Remarkably, mice immunized with MAP-flu displayed little to no weight loss while mice receiving Alum-flu lost on average 5% of their body weight, consistent with a less symptomatic infection and superior protection in MAP-flu vaccinated mice when compared to Alum-flu. Overall, these data demonstrate that MAP-flu immunization can elicit protection against lethal influenza infection.

## Conclusion

In this study, we demonstrate that VaxMAP can function as a robust immunization platform for either protein-based or inactivated pathogen-based immunogens, two traditional vaccine types commonly used to vaccinate patients against a variety of viral and bacterial pathogens. We demonstrate that VaxMAP improved immune responses to a classical immunological adjuvant and provided enhanced protection against symptomatic infection in a vaccination model against influenza virus when compared to Alum. Further, by simply changing the weight % of PEG, without adding cytokines, growth factors, or PAMPs, DAMPs or other adjuvants, we were able to fine-tune immune responses leading to improved antigen specific antibody responses, without affecting polarization of immunity from type 2 to type 1 immune responses. Immunization with VaxMAP drives activation of CD4^+^ T cells and B cells, as well as their differentiation to the adaptive Tfh and GC B cells, respectively. These immune cells are critical regulators of germinal center responses, which lead to the generation of long-lived memory B cells and plasma cells, required for extended protection against pathogens. Finally, we show that immunizing mice with 7.5% VaxMAP loaded with inactivated influenza virus completely protected mice against a lethal influenza viral challenge 60 days after vaccination, demonstrating VaxMAP-based immunizations can serve as functional vaccines.

Immunologically instructive biomaterials have shown potential as novel vaccination platforms. Most vaccine adjuvant platforms, including the promising mesoporous silica nanoparticle platform and the recently FDA-approved Matrix M platform, are based on nanoparticles that do not have cell-scale porosity^[53–55]^. Biomaterials with macroscale porosity that allow rapid cellular infiltration are being recognized for their potential to modulate immune responses in tissue regeneration, cancer immunotherapy, and immune-tolerance applications, but their ability to generate immune responses in traditional vaccine settings remains incompletely explored. Super et al. recently demonstrated the ciVAX platform using mesoporous silica particles coupled with the TLR4 agonist lipid A and a Fc-mannose-binding lectin to bind bacterial components to protect mice from bacterial infections^[34]^. This platform demonstrated the ability to promote antibody production, decrease bacterial burden, and prevent mortality, but whether the protective effects could be produced by the biomaterial itself, or the numerous factors added to the biomaterial is unknown. Our work provides evidence that a biomaterial alone can provide the necessary signals to induce Tfh and GC B cells, protective antibodies, and protection against lethal infection. In fact, the biomaterial-induced immune response is tunable and we demonstrate that altering material crosslinking density, which enhances material stiffness, as a novel means to enhance the adjuvant effects of a biomaterial platform to generate more robust antibody responses and improved protection against a viral infection when compared to traditional platforms. This enhanced immune response occurs without altering the polarization of the immune response, which could be desired for neutralizing toxins or viral attachment proteins., Future studies may uncover whether modulation of other material cues, such as adhesion peptide concentration, adhesion peptide sequence tuned to recruit specific immune cells^[56]^ or biomaterial composition alter response to type 1 immune polarization rather than type 2 immune polarization to provide protection against different types of pathogens^[57]^. Thus, our study provides support for the use of a modular injectable biomaterial to directly induce protective immune responses for vaccine applications.

### Experimental Section

#### Microfluidic Device Fabrication

Droplet generating microfluidic devices were fabricated by soft lithography as previously described^[35]^. Briefly, master molds were fabricated on silicon wafers (University wafer) using two-layer photolithography with KMPR 1050 photoresist (Microchem Corp). The height for the droplet formation channel was 50 μm, and the height for the collection channel was 150 μm. Devices were molded from the masters using poly(dimethyl)siloxane (PDMS) (Sylgard 184 kit, Dow Corning). The base and crosslinker were mixed at a 10:1 mass ratio, poured over the mold and degassed before curing overnight at 65 °C. Channels were sealed by treating the PDMS mold and a glass microscope slide (VWR) with oxygen plasma (Plasma Cleaner, Harrick Plasma) at 500 mTorr and 80 W for 30 s. Thereafter, the channels were functionalized by injecting 100 μL of Aquapel (88625-47100, Aquapel) and reacting for 30 s until washed by Novec 7500 (9802122937, 3M). The channels were dried by air suction and kept in the oven at 65 °C until used.

#### Microgel Production

Monodisperse microgels were produced as follows. Two aqueous solutions were prepared: i) 4 Arm-PEG VS (PTE-200VS, JenKem) at 10% and 15% (w/v) in 0.3 M triethyloamine (TEOA) pH8.5, pre-reacted with K-peptide (Ac-FKGGERCG-NH2), Q-peptide (Ac-NQEQVSPLGGERCG-NH2) and with RGD peptide (Ac-RGDSPGERCG-NH2) with 1mg/mL NP-ova and ii) an di-cysteine modified matrix metallo-protease sensitive peptide (MMP) (Ac-GCRDGPQGIWGQDRCG-NH2) (Genscript). These pre-gel solutions were sterile-filtered through a 0.2 μm polyethersulfone (PES) membrane in a luer-lok syringe filter, injected into the microfluidic device and pinched off by oil phase (0.1% Pico-Surf in Novec 7500, SF-000149, Sphere Fluidics) **(Figure S1, Supporting Information)**. The flow rate for aqueous solutions was 4–6 μL min^−1^ and for oil solutions was 30–100 μL min^−1^ to fine-tune the size of droplets. Gels were collected from the device into a tube in oil phase, incubated overnight at room temperature in dark. Microgels in oil phase were vortexed with 20% 1H,1H,2H,2H-Perfluoro-1-octanol (PFO) (370 533-25G, Sigma-Aldrich) in Novec 7500 for 10 s. Microgels were then mixed with 1:1 mixture of HEPES buffer (100 × 10^−3^m HEPES, 40 × 10^−3^m NaCl pH 7.4) and hexane followed by centrifugation at 10,000 rpm to separate microgels from oil for five times. Microgels were incubated in sterile-filtered 70% ethanol solution at 4 °C at least overnight for sterilization. Before in vivo or in vitro experiments, microgels were washed with a HEPES buffer with 10 × 10^−3^m CaCl_2_ for five times.

#### Annealing of Microgels

Equal volumes of two microgel solutions were incubated in HEPES-buffered saline (pH 7.4) containing FXIII (10 U mL^−1^) or thrombin (2 U mL^−1^) respectively at 4 °C overnight. The two solutions were centrifuged at 10,000 rpm for 5 min and supernatants were removed to concentrate the microgels. These concentrated solutions were thoroughly mixed with each other by pipetting up and down, pipetted into a desired location and kept at 37 °C for 90 min to anneal the microgels into a MAP scaffold.

#### Rheology Techniques for Measuring the Storage Modulus of MAP Blocks

The storage modulus of an 8 mm disc gel was measured using an Anton paar physica mcr 301 Rheometer. 40 μL of pre-gel solutions (20 μL of PEG with peptides, 20 μL of crosslinker) were pipetted onto sterile siliconized (Sigmacote; SL2-25ML, Sigma-Aldrich) slide glass, covered with another glass with 1 mm spacer and incubated at 37 °C for 2 h. Disc gels were swollen to equilibrium in PBS overnight before being measured. An amplitude sweep (0.01–10% strain) was performed to find the linear amplitude range for each. An amplitude within the linear range was chosen to run a frequency sweep (0.5–5 Hz). At least four disc-gels were measured for each condition.

#### Void Fraction of MAP Scaffolds

Fully swollen and equilibrated MAP building blocks (20 μL) were activated by with 5 U mL^−1^ FXIIIa (Sigma) and 1 U mL^−1^ thrombin, and the mixture was pipetted into a 3 mm diameter PDMS well on a glass coverslip and annealed in a humidified incubator at 37 °C for 1.5 h to form porous MAP scaffolds. Thereafter, the scaffolds were placed into HEPES buffer (pH 7.4) with 70 kDa dextran-FITC (FD70S-100MG, Sigma-Aldrich) overnight to reach equilibrium. Samples were 3D imaged using a Leica TCS SP8 confocal microscope with 10× objective.

#### Hydraulic Conductivity Measurement in the Scaffold

A custom-designed device was designed using Autodesk Inventor 3D CAD software and printed in Watershed XC 11 122 Normal-Resolution Stereolithography built in 0.004” layers from Proto Labs, Inc. **(Figure S2, Supporting Information)**. For the MAP scaffold, 25 μL of microgel building blocks (5 or 7.5 wt% crosslinked with MMP-cleavable dithiol) was casted on top of a 5 μm pore size cellulose membrane (SMWP01300, Fisher Scientific) in the bottom plane of the device and annealed followed by the overnight incubation in PBS. For the nonporous scaffold, 10 μL of pre-gel solution was casted on top of the membrane in the device and incubated at 37°C for 2 h followed by the overnight incubation in PBS. Then 1 mL of PBS with blue food dye was injected into the device and the permeated volume over time was measured. The hydraulic conductivity was calculated based on Darcy’s law.

#### Antigen release from MAP gel

FITC conjugated Ovalbumin (Invitrogen) as surrogates for hydrogel degradation and antigen release from hydrogels after a 3-day incubation in PBS and collagenase solution at the different concentrations.

#### Bone Marrow Derived Macrophage and Dendritic Cell Culture in MAP scaffold

Bone marrow was harvested from C57BL/6J (B6) mice. Macrophages were differentiated using CMG media. Dendritic Cells were differentiated using 20 ng/mL Recombinant Murine GM-CSF (Peprotech) and 10 ng/mL IL-4 (Peprotech). Cells were collected at Day 6 for in vitro assays. Collected cells were mix with MAP gel scaffold and co-cultured for a week. Before microscope observation, cells were stained by Hoechst 33342 Solution (Thermo Fisher Scientific).

#### Mice

Mice were housed in pathogen-free conditions at the Rutgers New Jersey Medical School or UCLA. C57BL/6J (B6) were purchased from NCI managed colony at Charles River or Jackson Laboratories. All animals were ages 6–12 weeks during the course of the study, with approval for procedures given by the Institutional Animal Care of Rutgers New Jersey Medical School or UCLA. Both male and female mice were used in these studies.

#### VaxMAP Injection

For VaxMAP injection the same volume of HEPES buffer is mixed with MAP microgels. Excess supernatant is removed as much as possible. A Factor XIII and thrombin solution in the ratio of 6U Factor XIII/1.2 U thrombin solution is made in 200 μL HEPES buffer and added to MAP microgels in the ratio of 1mL gel to 200μL Factor XIII/Thrombin. Final MAP microgel solution is loaded into a Hamilton GasTight syringe and injected s.c. at the base of the tail with a 25G needle.

#### Flow Cytometry and Cell Sorting

Tissues were homogenized by crushing with the head of a 1-ml syringe in a Petri dish followed by straining through a 40-μm nylon filter. Ammonium–chloride–potassium buffer was used for red blood cell lysis. To determine cell counts, count beads were used at a concentration of 10,000 beads/10L and added directly to samples following staining. Cell count is normalized by dividing input bead count by cytometer bead count, and multiplied by the dilution factor. Antibodies used for flow cytometry staining are listed in Table S1. Staining for CXCR5 was performed at room temperature (25°C) with 30 min of incubation. Stained and rinsed cells were analyzed using a multilaser cytometer (LSRII; BD Biosciences or Attune; Invitrogen).

#### Influenza Inactivation and Infections

1mL stocks of live PR8 Influenza were inactivated by incubating with 1μL Beta-propiolactone overnight at 4 degrees. For infections, mice were anesthetized with 87.5mg/kg Ketamine/ 12.5 mg/kg Xylazine cocktail in 200μL of PBS prior to infection. Mice were infected intranasally with 30μL of 0.5LD_50_ influenza PR8 in PBS using a p200 pipette. Animals were sacrificed at indicated time points p.i. or if they lost 30% of their initial weight and harvested organs were processed for flow cytometry.

#### ELISA Assays

Anti-NP and anti-HA antibodies in mouse sera were measured on Nunc PolySorp 96 well plates (Thermo Fisher) coated with NP-9 or NP-27 protein (Biosearch Technologies) or recombinant HA1 (Sino biologics) in carbonate buffer (Sigma). Alkaline phosphatase-conjugated anti-mouse IgG antibodies (Southern Biotech) and phosphatase substrate (Sigma) were used for detection. ODs were read at 405 nm on a SpectraMax Microplate Reader (Molecular Devices).

#### Statistical Analysis

Data were analyzed using Student’s *t*-test, Mann-Whitney unpaired *t*-test, and Pearson correlation coefficient with GraphPad Prism 8 Software.

## Supporting Information

Supporting Information is available online.

## Acknowledgements

We acknowledge support from the UCLA W. M. Keck Foundation COVID 19 Research Award Program (D.D., P.O.S.), the Presidential Early Career Award for Scientists and Engineers (N00014-16-1-2997) (D.D.), NIH R01-AR079470-01 (P.O.S. and D.D), and LEO Foundation LF-OC-21-000688 (P.O.S.) for funding. O.Q.A. was supported by NIH 2 T32AI125185 J.S.W. and laboratory was supported in part by National Institutes of Health (NIH) grant R01 AR073912. Device mold fabrication was completed using equipment provided by the Integrated Systems Nanofabrication Cleanroom at the California NanoSystems Institute at the University of California, Los Angeles.

