## Supporting Information for "Improved humoral immunity and protection against influenza virus infection with a 3D porous biomaterial vaccine"

K.M. Kelly-Scumpia
Division of Cardiology, Department of Medicine David Geffen School of Medicine University of California, Los Angeles, Los Angeles, CA 90095, USA

P.O. Scumpia
Division of Dermatology, Department of Medicine David Geffen School of Medicine University of California, Los Angeles, Los Angeles, CA 90095, USA


M.M. Archang,
MSTP Program, David Geffen School of Medicine, University of California Los Angeles, Los Angeles, CA 90095, USA

P.O. Scumpia
Department of Dermatology VA Greater Los Angeles Healthcare System Los Angeles, CA 90073, USA

D. Di Carlo
Department of Mechanical and Aerospace Engineering University of California, Los Angeles Los Angeles, CA 90095, USA

D. Di Carlo
California Nano Systems Institute (CNSI) University of California, Los Angeles Los Angeles, CA 90095, USA

P.O. Scumpia, D. Di Carlo
Jonsson Comprehensive Cancer Center University of California, Los Angeles Los Angeles, CA 90095, USA

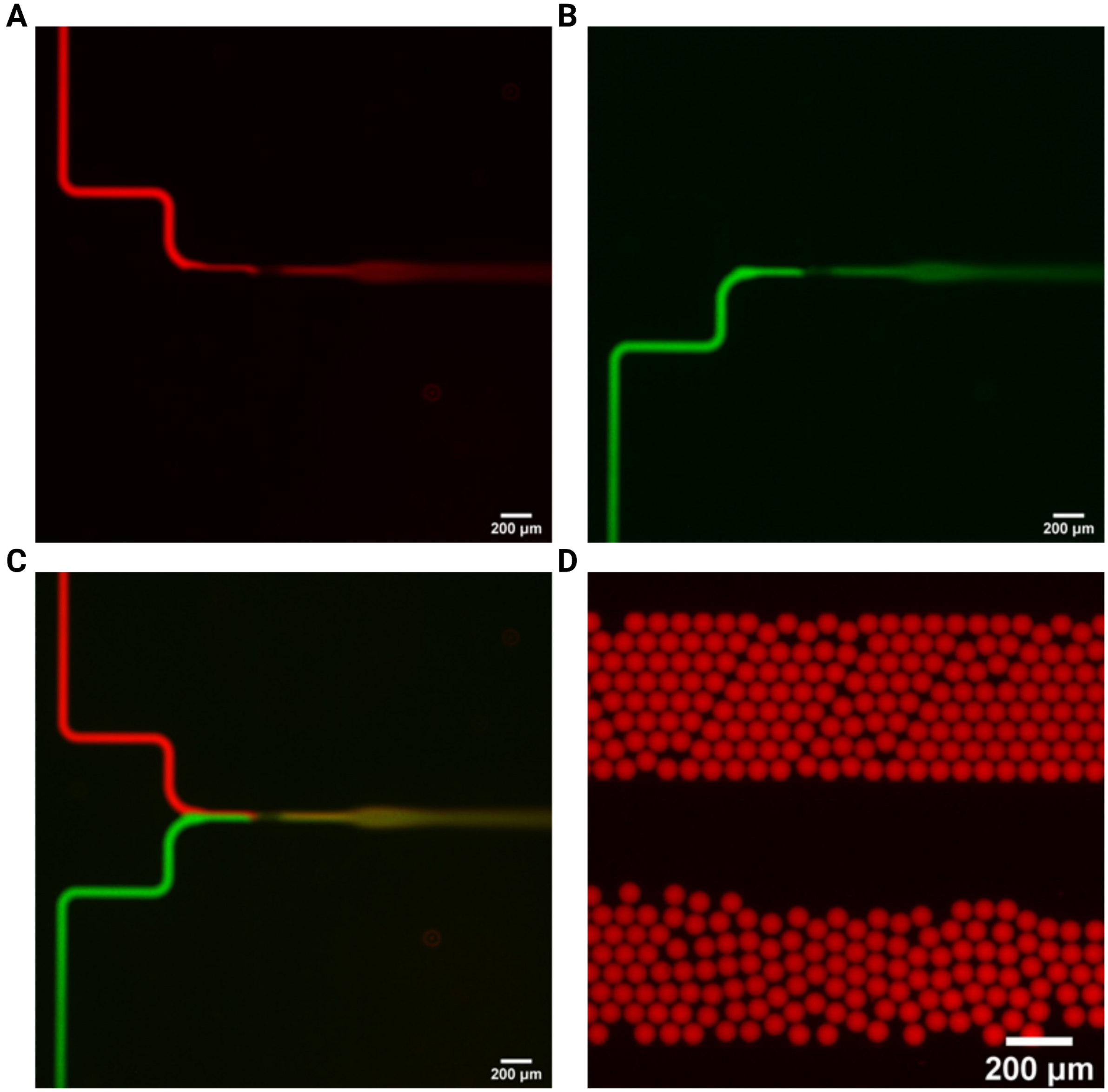


**Figure S1.** Microfluidics VaxMAP fabrication. Channels are highlighted with fluorescent dye solutions. Homogeneous droplets containing pregel solution and crosslinker form at a flow focusing junction of the microfluidic channel. A) The aqueous inlet channel with fluorescently tagged OVA (green) contains 4-arm PEG-vinylsulfone pregel solution. B) The second aqueous channel contains AlexaFluor 546-maleimide (red) with MMP-sensitive cross linker solution. C) Merged channel image. D) Fluorescence images of droplets generated downstream. Scale bars are 200 μm.

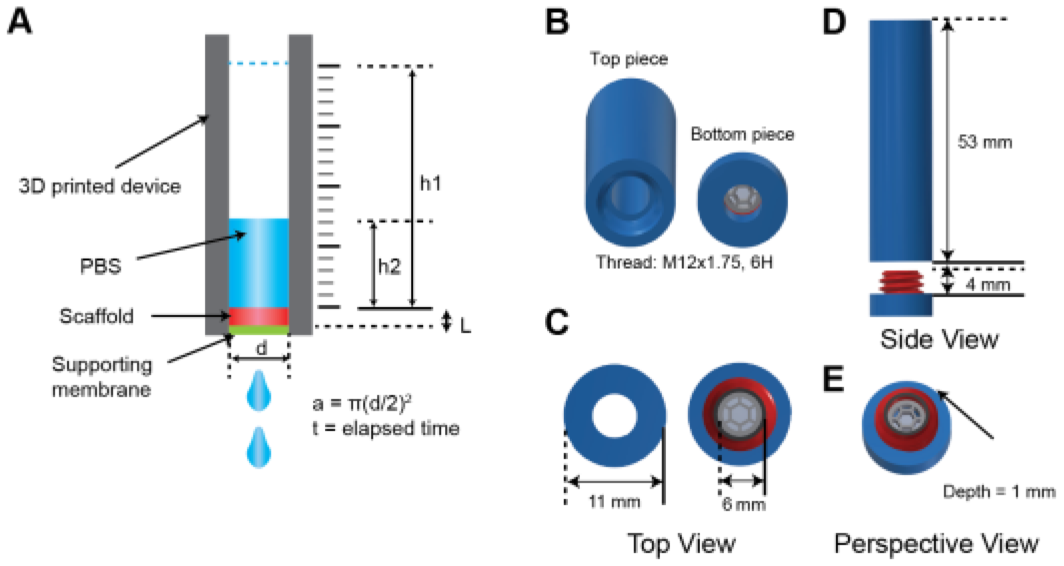


**Figure S2.** Hydraulic conductivity measurement apparatus. A) Schematic of the hydraulic conductivity measurement using a 3D-printed device. The initial height (h1) and final height (h2) of PBS that flows through the scaffold over an elapsed time were recorded to calculate the overall volumetric flow rate and the conductivity. B) Two components of the 3D-printed device viewed from C) top, D) side and E) perspective.

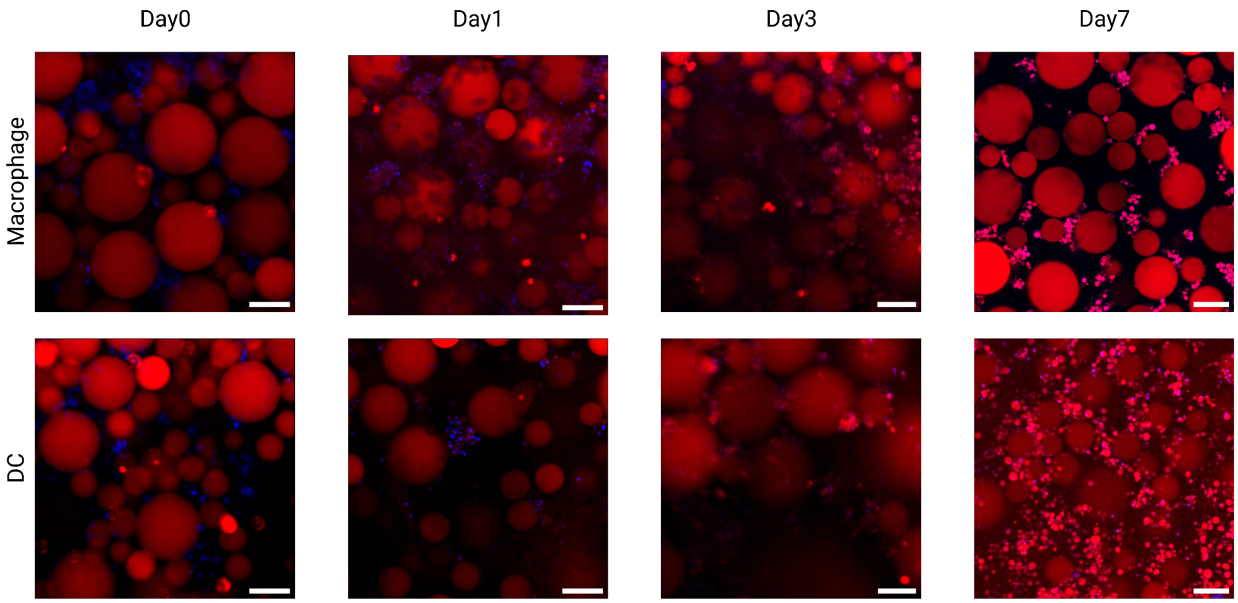


**Figure S3.** In vitro culture of mouse bone marrow derived macrophages and dendritic cells (DCs). Cells are cultured within 3D MAP scaffolds without antigen over 7 days. As the incubation period increased, MAP-derived fluorescence signal (Alexa Fluor 555-maleimide) accumulated in the cells. Blue: DAPI stain, Red: Alexa Fluor 555. Scale bars are 100 μm.

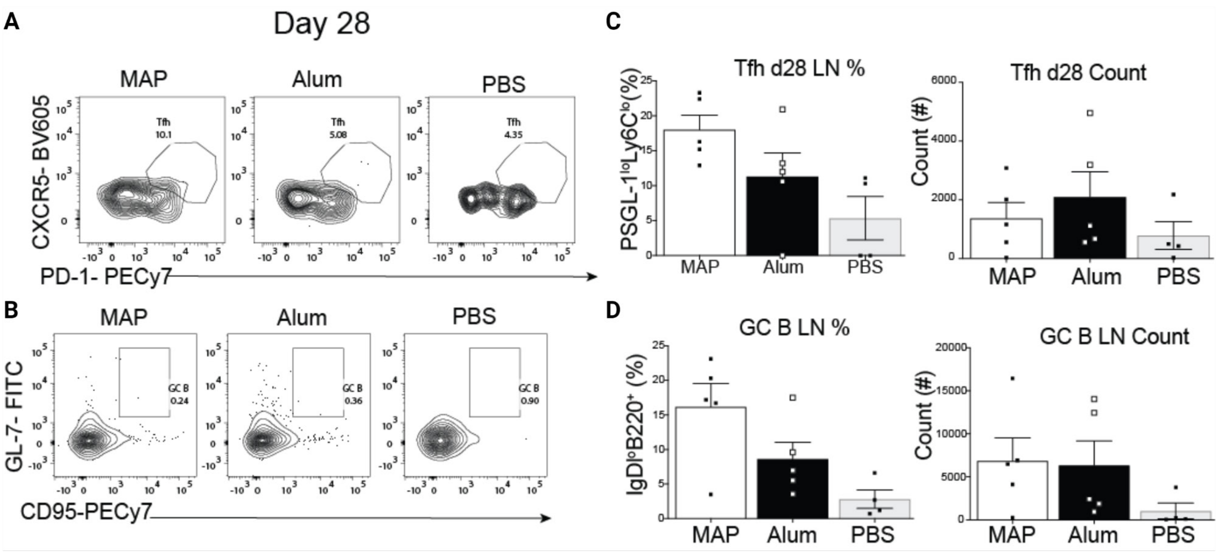


**Figure S4**. Tfh and GC B cell responses at day 28 post immunization from 5% VaxMAP injection. A-B) Representative FACS plots, percentages and counts of Tfh cells. C-D) Representative FACS plots, percentages and counts of GC B cells.

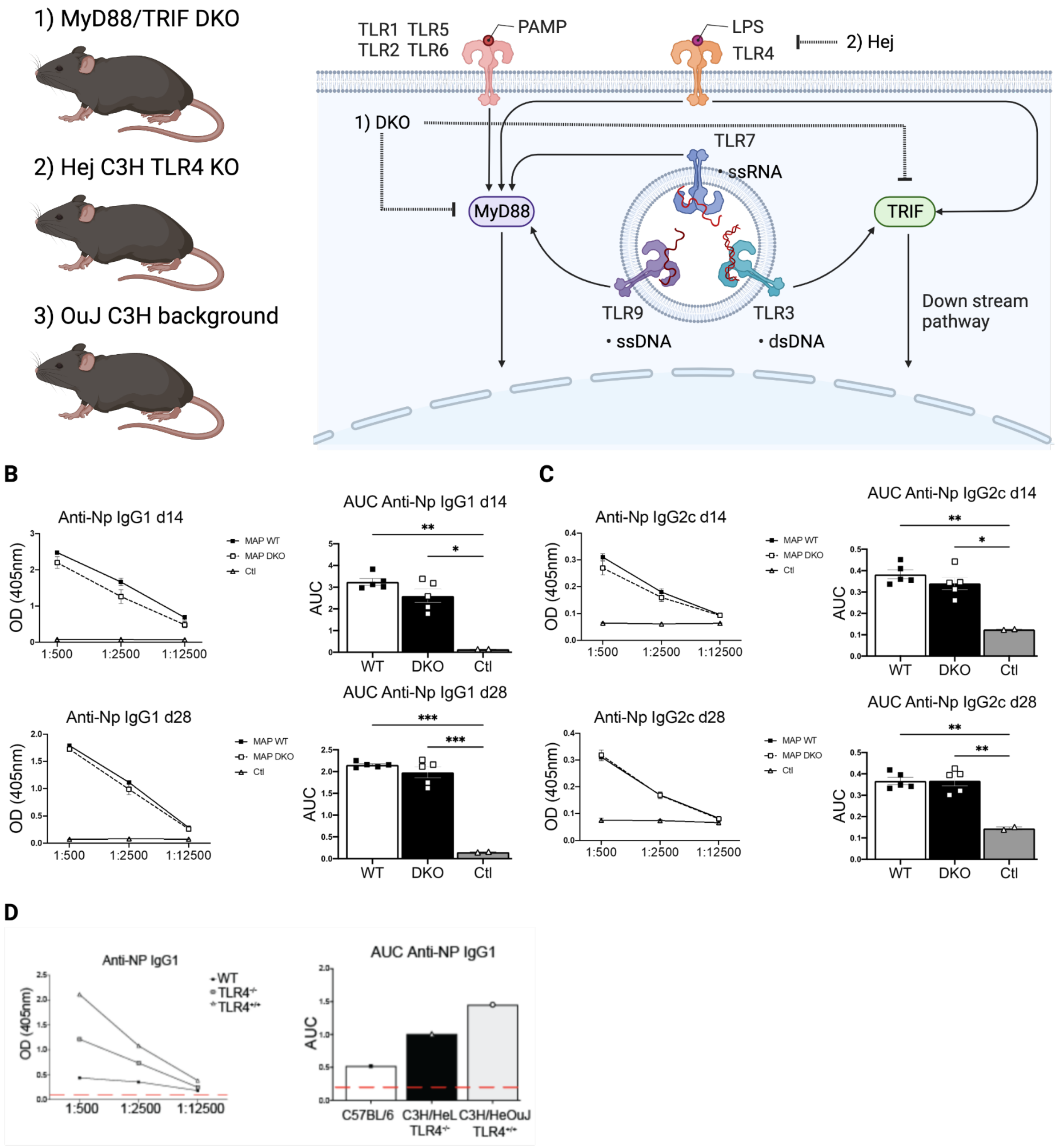


**Figure S5.** MyD88/TRIF DKO. A) Schematic of signaling pathway of importance in innate immunity. B) Optical Density and Area Under the Curve of anti-NP IgG1 antibodies at day 14 (top) or 28 (bottom) post immunization. C, D) Optical Density and Area Under the Curve of anti-NP IgG2c antibodies at day 14 (top) or 28 (bottom) post immunization.

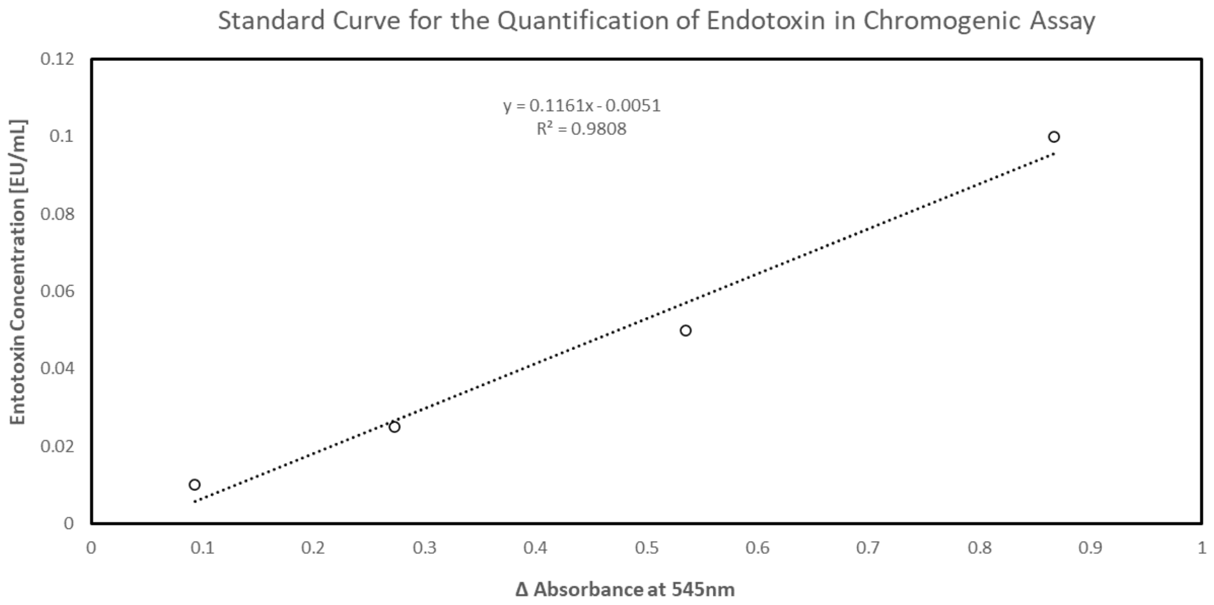


**Figure S6.** Endotoxin assay. VaxMAP is tested for concentration of endotoxin using an assay kit (Genscript ToxinSensor™ Chromogenic LAL Endotoxin Assay Kit). Fabricated VaxMAPs are mixed with ToxinSensor reagents and absorbance at 545nm is measured to confirm endotoxin concentration was less than 0.01 EU/mL before animal testing. Standard curve is shown to calibrate threshold levels.

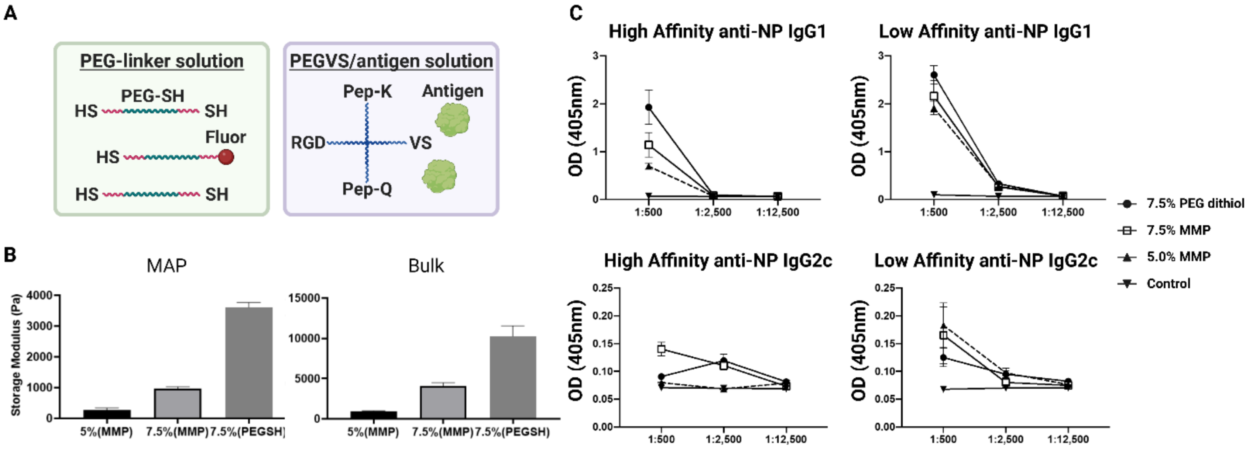


**Figure S7.** Effect of MAP gel formulation on high and low affinity antibody titers. A) MAP microgels produced with 4-arm PEG-vinylsulfone (PEG-VS) via thiol-ene reactions to encapsulate antigen in the dense gel mesh. PEG dithiol was used as a crosslinker to enable microparticle formulations with higher stiffness than MMP crosslinkers. B) Stiffness of PEG dithiol crosslinked gels formed in bulk and annealed MAP gel form. PEG dithiol crosslinking resulted in more than 2-fold increased stiffness than the 7.5% MMP crosslinked condition. C) Optical density of high affinity (NP-9) and low affinity (NP-27) anti-NP IgG1 and IgG2c antibodies at 14 days post immunization. PEG represents PEG dithiol crosslinked condition. PEG dithiol was crosslinked with 7.5% of PEG-VS. 7.5% and 5% MAP represented MMP crosslinked conditions.

**Table S1.** Antibodies Used for Flow Cytometry

| Antigen | Dilution | Clone | Fluorochrome | Source | Cat# |
|---|---|---|---|---|---|
| CD4 | 1:200 | RM4-5 | APC | Biolegend | 100516 |
| CD4 | 1:200 | RM4-5 | AF700 | Biolegend | |
| CD44 | 1:200 | IM7 | APC Cy7 | Biolegend | 103028 |
| B220 | 1:200 | RA3-6B2 | BV605 | Biolegend | 103244 |
| IgD | 1:200 | 11-26c.2a | BV421 | Biolegend | 405225 |
| GL-7 | 1:200 | GL-7 | Alexa488 | Biolegend | 144612 |
| CD95 | 1:200 | Jo2 | PE Cy7 | BD Pharmagen | 557653 |
| PSGL-1 | 1:1000 | 2PH1 | Pacific Blue | BD Biosciences | 562807 |
| Ly6C | 1:400 | HK1.4 | BV510 | Biolegend | 128033 |
| PD-1 | 1:200 | 29F.1A12 | PE-Cy7 | Biolegend | 135216 |

| | | | | | |
|---|---|---|---|---|---|
| CXCR5 | 1:100 | L138D7 | BV605 | Biolegend | 145513 |

**Table S2.** Chemicals and Peptides

| Reagent or Resource | Source | Cat # |
|---|---|---|
| Phosphate Buffered Saline (PBS) | Sigma | 806544 |
| RPMI-1640 | Corning | **10-040-CM** |
| Fetal Bovine Serum | VWR | 89510-186 |
